# cFos ensembles in the dentate gyrus rapidly segregate over time and do not form a stable map of space

**DOI:** 10.1101/2020.08.29.273391

**Authors:** Paul J. Lamothe-Molina, Andreas Franzelin, Lennart Beck, Dong Li, Lea Auksutat, Tim Fieblinger, Laura Laprell, Joachim Alhbeck, Christine E. Gee, Matthias Kneussel, Andreas K. Engel, Claus C. Hilgetag, Fabio Morellini, Thomas G. Oertner

**Author notes:** equal contribution (Paul J. Lamothe-Molina, Andreas Franzelin). these authors co-supervised the study (Fabio Morellini, Thomas G. Oertner).

## Abstract

Transgenic cFos reporter mice are used to identify and manipulate neurons that store contextual information during fear learning. It is not clear, however, how spatial information acquired over several training days is integrated in the hippocampus. Using a water maze task, we observed that cFos expression patterns in the dentate gyrus are temporally unstable and shift daily. Surprisingly, cFos patterns did not get more stable with increasing spatial memory precision. Despite the fact that cFos was no longer expressed, optogenetic inhibition of neurons that expressed cFos on the first training day affected performance days later. Triggered by training, ΔFosB accumulates and provides a negative feedback mechanism that makes the cFos ensemble in the dentate gyrus dependent on the history of activity. Shifting cFos expression to a different set of granule cells every day may aid the formation of episodic memories.

## Introduction

Accurate memory of past events has a high adaptive value for animals, as all decision making is based on past experience. The hippocampus is well-known for its processing of spatial information ^1,2^ but it also processes other types of sensory input and is capable of grouping events together in time to form episodic memories ^3–5^. Our understanding of how time is represented in the hippocampus is incomplete. The expression of many genes affecting synaptic plasticity undergoes pronounced circadian oscillations, changing the rules of synaptic plasticity depending on the time of day ^6^. Theoretical and empirical work suggests that the dentate gyrus (DG) actively reduces the overlap between activity patterns from the entorhinal cortex, a process dubbed pattern separation ^7–9^. In vivo electrophysiology and calcium imaging experiments revealed that the majority of the granule cells (GCs) in the DG are silent, and of the active cells, just a small fraction show spatial tuning (place cells) ^10–12^. Spatially tuned GCs provide a stable, albeit coarse representation of the global environment across time ^10,13–15^. These findings highlight an apparent design conflict: while a perfect ‘episode encoder’ should avoid using the same neurons on consecutive days, accurate place coding is thought to require stable place cells that signal the animal’s position in a given environment. In view of this conundrum, we set out to study the impact of time and space on GCs in the dorsal DG.

To monitor and manipulate neuronal activity in freely behaving animals, reporter mice have been developed that use the immediate-early gene cFos ^16^ to drive the expression of fluorescent proteins and optogenetic actuators ^17^. Fear conditioning experiments with activity-dependent expression of optogenetic silencing tools suggest that recall of a fearful episode requires reactivation of the original encoding ensemble ^18–20^. Vice versa, artificial reactivation of the encoding ensemble has been shown to reinstate a fearful state ^19,21,22^, suggesting that memories can be activated by specific subsets of hippocampal neurons. When freezing is used as a proxy to assess the emotional state of the animal, it is difficult to determine whether, during the optogenetic reactivation, the animal recalls the fear-inducing episode (“engram”) or simply feels fear. The reactivated ensemble may form an internal representation of the external world (cognitive map theory ^23^) or trigger the pattern of cortical activity that was active in the previous experience (indexing theory ^24^).

We investigated the temporal stability of cFos ensembles and their relation to spatial learning and the formation of cognitive maps. We used cFos-dependent tagging to assess how cFos-ensemble overlap changes over training days in the Morris water maze (WM). Even in expert mice, cFos expression patterns in DG changed from day to day. In spite of these changing expression patterns, optogenetic inhibition of cFos-tagged GCs impaired navigation 5 days after tagging, suggesting that the absence of cFos expression in these GCs did not imply absence of activity. We show in vivo and in vitro that after a period of cFos expression, cFos^+^ GCs accumulate ΔFosB, a long-lived splice variant of FosB. As this splice variant inhibits cFos expression ^25^, it provides a potential mechanism for the daily shift of cFos ensembles that occurs even when mice are exposed to the same environment. The changing pattern of cFos expression in DG could be the basis of an episodic memory system that write-protects synapses on the most recently used subset of GCs during the following days.

## Results

### Spatial learning and cFos ensemble overlap in the dentate gyrus: novice vs. expert mice

Although cFos expression is considered a proxy of neuronal activity ^26–28^, the relationships between action potential firing, immediate early gene expression, and synaptic long-term plasticity are not obvious ^29^. Place cells fire bursts of action potentials when a mouse is in a specific location in space, and place cell consistency is reportedly high in dentate gyrus (DG) ^10^. We chose a spatial learning paradigm, the Morris water maze (WM), to investigate cFos expression patterns in DG during learning. TetTag mice were trained to find a hidden platform in the WM (Fig. 1a-c). We labeled cFos ensembles from two consecutive days (Fig. 1d). The window for labeling the first cFos ensemble was opened by taking the mouse off doxycycline (Dox), allowing for long-term expression of the fluorescent protein mKate2 fused to an opsin for membrane labeling (cFos-tagged, Fig. 1d). A short half-life version of the green fluorescent protein (shEGFP) works as an in-built cFos reporter in the TetTag mouse to label the second cFos^+^ ensemble. In calibration experiments, we observed slightly faster temporal dynamics of the native cFos protein than of the shEGFP reporter, yet nearly all shEGFP-expressing neurons also expressed cFos (Supplementary Fig. 1). The highest correlation between native cFos and shEGFP was obtained 2-4 h after stimulation, we therefore chose 3 h after the last training trial as the timepoint to sacrifice mice and fix the brains. The first cFos-tagged ensemble was revealed by immunofluorescence staining against mKate2 and the second cFos-expressing ensemble was revealed by staining against the shEGFP (cFos^+^). The ensemble sizes (proportion of GCs cFos-tagged and cFos^+^, Supplementary Fig. 2) were used to calculate the amount of overlap (GCs that are both cFos-tagged and cFos^+^) expected due to chance for each sample and compared to the actual overlap (number of cFos-tagged cFos^+^ GCs).

**Figure 1.**
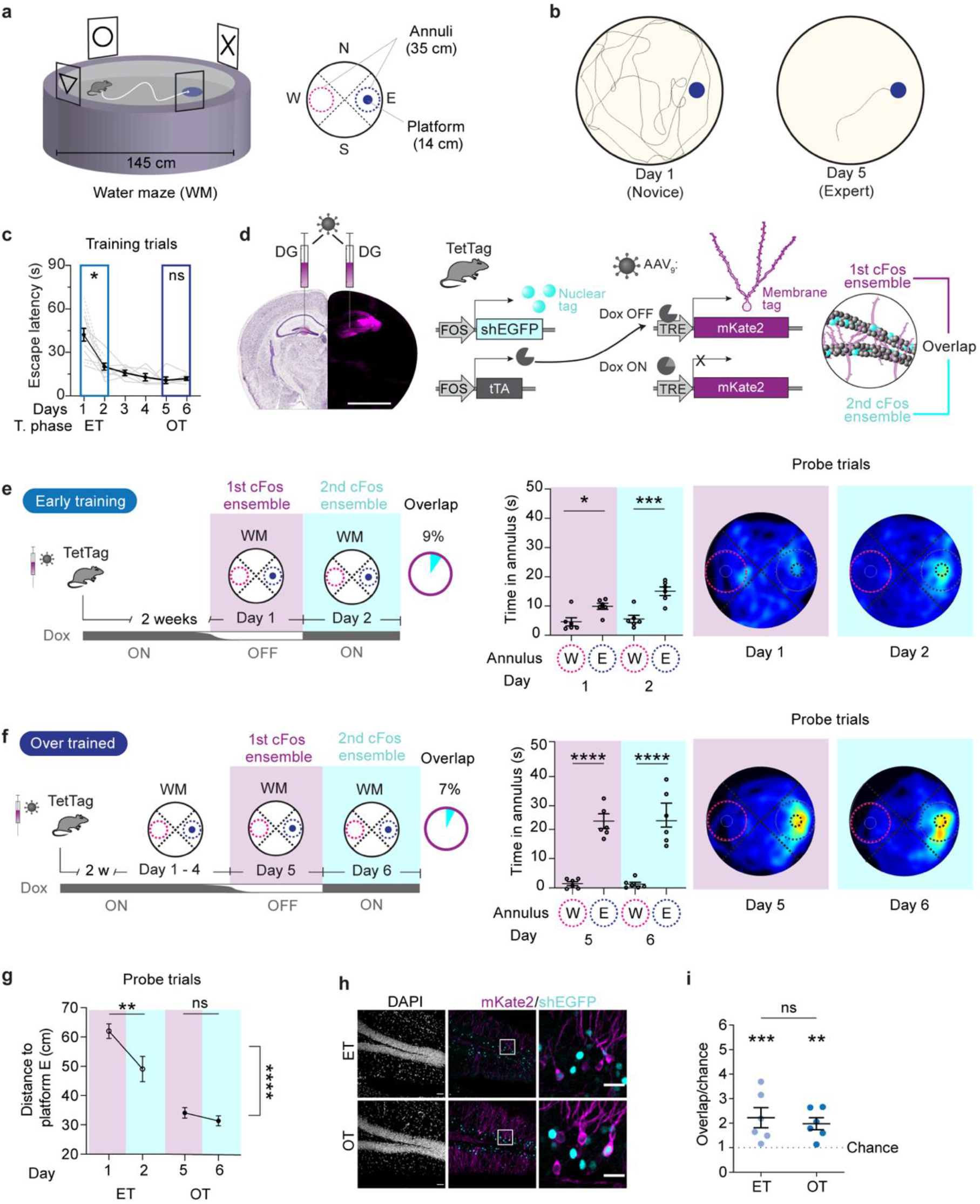
Behavioral performance in a spatial memory task is not reflected by cFos ensemble overlap in the DG. **a,** Hidden platform version of the WM. A 14 cm platform was placed in the east quadrant (E) 1 cm under the surface in a 145 cm diameter tank. In probe trials, time spent in 35 cm dia. annuli centered on platform E and in the opposite quadrant (W) was measured. **b,** Exemplary swim paths from days 1 & 5 of training. **c,** Escape latencies (s) over training days (black line represents average of all mice in both groups n = 12, dashed gray lines: ET group; solid gray lines: OT group). Mice improve rapidly from day 1 to 2 (* p = 0.02) but plateau after day 5 (ns p > 0.99), Sidak correction after mixed model ANOVA. **d,** In TetTag mice, the cFos promoter drives the expression of two transgenes: 1) tetracycline transactivator (tTA) and 2) a short half-life, nuclear-localized, enhanced green fluorescent protein (shEGFP). In the absence of doxycycline (OFF Dox), tTA binds to the tTA-response element (TRE) of a virally delivered construct, resulting in permanent cFos-tagging of neurons (1^st^ ensemble, mKate2, magenta). Nuclear fluorescence identifies neurons that were cFos^+^ in the hours before the animal was sacrificed (2^nd^ ensemble, shEGFP, cyan). Overlap is the % of mKate2^+^ (cFos-tagged) neurons expressing shEGFP on the following day (cFos^+^). **e,** Early training group mice (ET, n = 6 mice) were trained OFF Dox on day 1. On day 2, mice were trained ON Dox and sacrificed after the last trial. Spatial reference memory is shown by the spent time in annuli (middle) and as a heatmap (right) during probe trials (Time in annuli Day 1: * p = 0.03, Day 2: ** p = 0.001. Sidak correction after a two-way-ANOVA). **f,** Over-trained mice (OT, n = 6 mice) were trained OFF Dox on day 5. On day 6, mice were trained ON Dox and sacrificed after the last trial. Spatial reference memory is shown by the spent time in annuli (middle) and as a heatmap (right) during probe trials (Time in annuli. Day 5 **** p < 0.0001, day 6: **** p < 0.0001. Sidak correction after a two-way-ANOVA). **g,** Mean distance to platform E during the probe trials. Spatial search accuracy was significantly higher in the OT than the ET group (one-way-ANOVA, Group effect: **** p < 0.0001). ET mice improved their spatial accuracy between tagging days while OT mice did not (ET group: day 1 vs. 2, ** p = 0.004; OT group: day 5 vs. 6, ns p = 0.65. Sidak correction after a two-way-ANOVA). **h,** cFos overlap analysis. Scale bar: 25 μm. Both mKate2 and shEGFP immunoreactive cells (middle & right) were counted inside the GC layer (DAPI, left). **i,** cFos overlap was significantly higher than expected by chance in both groups (Observed vs. Expected (chance) overlap: ET group: *** p = 0.0008, OT group: ** p = 0.003. Sidak correction after two-way-ANOVA) but not similar in both groups (Overlap/chance ET vs. OT group: ns p = 0.99, Sidak correction after one-way-ANOVA)

During probe trials, mice early in training (ET, n = 6 mice, Fig. 1b-c, e) were less precise and accurate while searching for the target but within 5 days became experts (over trained group, OT, n = 6 mice, Fig. 1 b-c, f). Not only the time spent searching in an annulus around the hidden platform increased (Fig. 1 e, f) but also distance to platform was significantly lower in the expert mice (Fig. 1 g). Our expectation was that, if the cFos expression is linked to activity of the neurons that are important for solving the WM task, there would be a high degree of overlap between the cFos^+^ and the cFos-tagged neurons, which were labeled 1 day earlier. We also expected overlap to increase with increasing search accuracy in the probe trials. Instead, we observed that very few GCs expressed both mKate2 (cFos-tagged) and shEGFP (cFos^+^) and that both the overlap (ET 9% and OT 7% Fig. 1e, f) and the ensemble sizes were similar in novices and expert mice (Supplementary Fig. 2). In both ET and OT mice, the observed overlap was, however, significantly higher than expected due to chance (Fig. 1h, I; Supplementary Fig. 2). Surprisingly, there was no difference between the novices and experts and, assuming that cFos indicates the active GCs, less than 90% of the neurons that expressed cFos the first day were re-activated the following day. Thus, in the DG cFos overlap in GCs on consecutive days of WM training is independent of spatial search accuracy.

### Reversal training vs novel environment

We next investigated how novelty affects the overlap of cFos ensembles. Would a new position of the escape platform (reversal training) decrease ensemble overlap? The first cFos ensemble was labeled in expert mice on day 5 of WM training (OFF Dox Fig. 2a). The following day, these mice were trained (ON Dox) with the platform on the opposite side of the tank (reversal training, RT, n = 6 mice). The training with a new platform position had no effect on cFos overlap (9% similar to ET and OT groups, Fig. 1 e, f), although it dramatically decreased search accuracy in test trials after reversal training (Fig. 2 a). The mice now searched equally in both E and W annuli.

**Figure 2.**
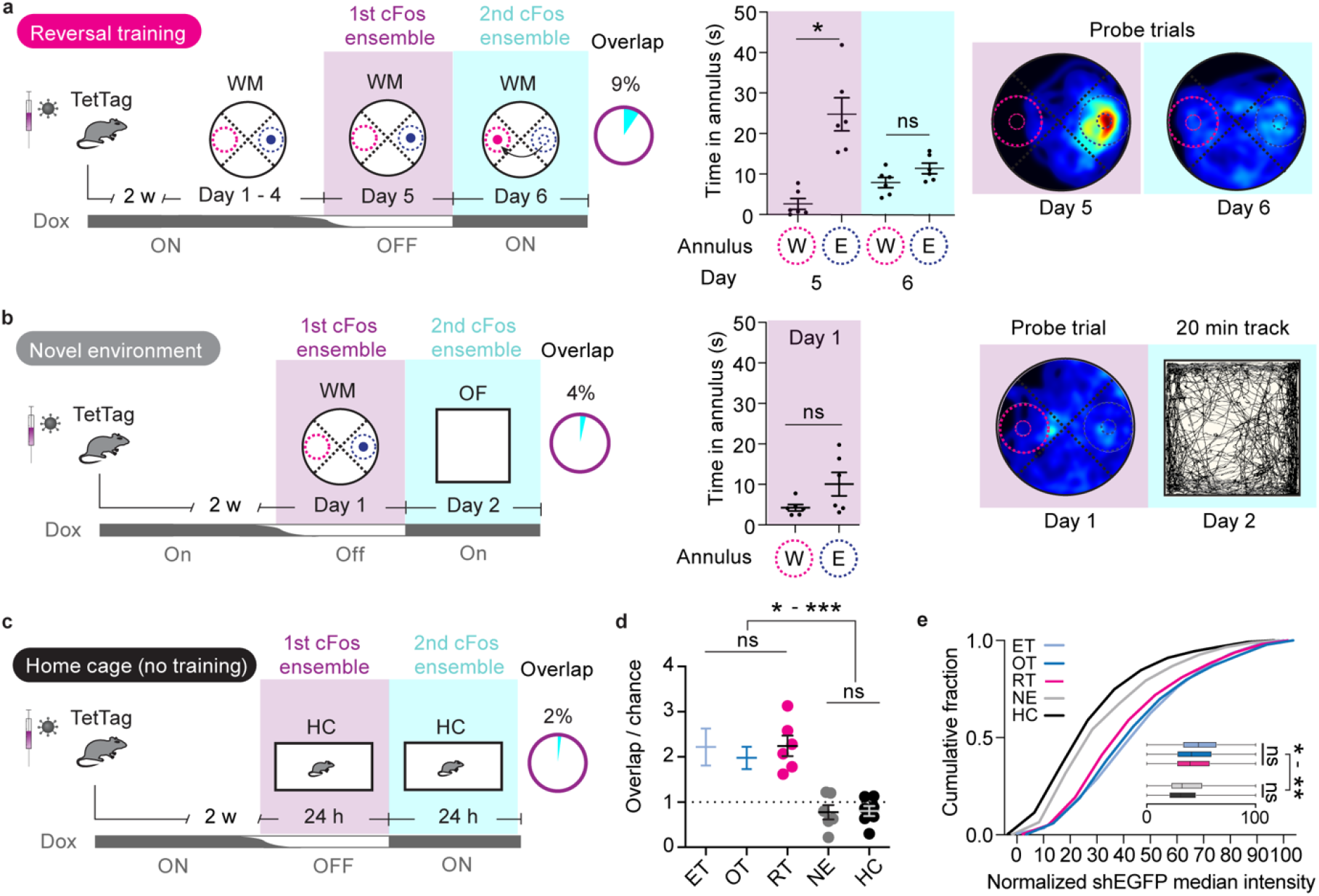
Effects of task and environment on cFos ensemble overlap. **a,** Reversal training group (n = 6). Mice were trained for 5 days with the platform in the east quadrant (E). On day 5, doxycycline was removed, and mice were further trained. On day 6, mice were trained ON Dox with the platform in the west quadrant (W). Spatial reference memory is shown by the spent time in annuli (middle) and as a heatmap (right) during probe trials. Despite the strong preference for E annulus on day 5 (*, p = 0.01), after only 1 reversal training session, mice searched equally around the new and the old platform position during a probe trial on day 6 (ns, p = 0.72). Sidak correction after two-way-ANOVA. **b,** Novel environment group (n = 6) was trained for 1 day OFF Dox in the WM. On day 2, mice explored an open field (OF) arena for 20 min Dox-ON. After 1 day of WM training, preference for the correct platform position was not yet significant (ns, p = 0.08, two-tailed t-test). Spatial reference memory is shown by the spent time in annuli (middle) and as a heatmap (right). Exemplary tracking during OF exploration (right) **c**, Control mice stayed in their home cage (HC group - no training, n = 7), had a 24 h OFF Dox period, and were sacrificed on day 2. **d,** cFos ensemble overlap/chance in the different experimental groups (ET, early training from Fig. 1i; OT, overtrained from Fig. 1i; RT, reversal training; NE, novel environment; HC, home cage). cFos overlap in the RT group is higher than expected by chance (***, p = 0.0006, see Supplementary Fig. 2). All WM-trained groups showed higher cFos overlap than the NE and HC groups (p values in supplementary table 1) but no difference between each other. **e**, cFos expression (normalized shEGFP median intensity) was significantly higher in mice that visited the WM than mice that visited an OF arena (NE group) or control mice (HC group) (p values in supplementary table 1).

To compare cFos overlap from different contexts, the first ensemble was cFos-tagged on day 1 of WM training. On the following day, mice explored an open field for 20 min (novel environment NE, n = 6 mice). The overlap in the NE group was 4%. We then compared overlap in mice that were never trained in the WM and kept in their home cages during days (HC group, n = 7). The cFos overlap was 2 % in the HC group. The cFos overlap of all groups trained in the WM (the ET, OT and RT mice) were not significantly different from each other but significantly higher than expected due to chance (Fig. 2d, Supplementary Fig. 2c). In contrast, in both the NE and HC groups, cFos overlap was at chance level and significantly lower than the WM groups. These findings were mirrored when we analyzed the intensity of shEGFP (Fig. 2e). Thus, WM training not only increased cFos overlap, which may be dependent on the scoring threshold, but also increased the amount of cFos expressed in the second ensemble (Fig. 2e, Supplementary Fig. 2). Thus, cFos is strongly driven in the WM, independently of training level (ET vs. OT) or difficulty of the task (OT vs RT). Home-cage labeled mice had the lowest number of cFos^+^ GCs (Supplementary Fig. 2), consistent with the concept that the hippocampus is highly engaged in spatial navigation tasks.

### cFos^+^ neurons in DG participate in spatial memory recall days later

Lesion ^30,31^ and optogenetic manipulation ^32–35^ experiments indicate that the DG plays a role in spatial memory acquisition and recall. However, these experiments did not address if those particular memories are encoded by a specific subset or ensemble of neurons in the DG (“engram cells”). Based on our finding that there is above-chance cFos ensemble overlap in mice that underwent WM training, we speculated that cFos^+^ cells in the DG may encode relevant information needed to solve the WM task. We employed a bidirectional optogenetic tool that can be used to inhibit or excite neurons with 473 and 594 nm light, respectively (BiPOLES ^36^). As we observed weak unspecific expression of mKate2 using the TREtight promoter (Supplementary Figure 3), we expressed BiPOLES under control of the non-leaky TRE3G promoter. BiPOLES functionality was confirmed in acute hippocampal slices prepared 24 h after cFos-tagging, in which BIPOLES-expressing GCs were targeted for patch-clamp recordings. As expected, 473 nm light prevented action potentials induced by current injection, while 594 nm light pulses repeated at 20 Hz reliably elicited single action potentials (Supplementary Figure 4). For in vivo experiments, mice were bilaterally injected in DG with AAV_PHP.eB_-TRE3G-BiPOLES-mKate2 and implanted with a custom-made tapered fiber implant ^37^ (Fig. 3a). To perform optogenetic experiments in the WM, we had to ensure that the weight of the optical fibers was compensated when mice were tethered to them via their implants. We achieved this by attaching a helium balloon to the optical fibers (Fig. 3a). Implanted mice were trained without being tethered and successfully learned the WM task (Fig. 3b). Attachment to the fiber optic cables did not affect swimming speed (Fig. 3c), nor did 473 or 594 nm light (Fig. 3d).

To investigate the importance of day 1 cFos^+^ neurons on memory recall, we silenced the BiPOLES-expressing neurons during the probe trials on days 2, 3, and 5 (Fig. 3f). Mice were placed on an elevated Atlantis platform that submerged after 30 s, forcing the mice to swim ^38^. After 60 s, a second Atlantis platform (in quadrant E) was elevated, and mice were directed towards it if they did not find it themselves. We analyzed the first half of the probe trials (30 s) to assess memory performance. Optogenetic inhibition did not affect the time spent in the target quadrant on days 2 and 3 when mice were novices and their spatial accuracy was still poor (Fig. 3g). On day 5, when mice were expert and spatial accuracy was high, optogenetic inhibition significantly decreased the time spent in the target quadrant (Fig. 3h, Supplementary Video 1). The mean distance to the target platform confirmed the results of the quadrant analysis, detecting a significant effect of optogenetic inhibition on days 3 and 5 (Fig. 3i). We confirmed that GCs still expressed BiPOLES even 9 days after tagging (Fig. 3j). These results suggest that the small number of cFos-tagged cells from the first training day (~2% of GCs) are important for successful memory recall on subsequent days. BiPOLES can also be used for selective reactivation of tagged cells (Supplementary Figure 4). However, optogenetic reactivation during day 6 probe trials had no significant effect on WM performance in expert mice (Supplementary Figure 6).

**Figure 3.**
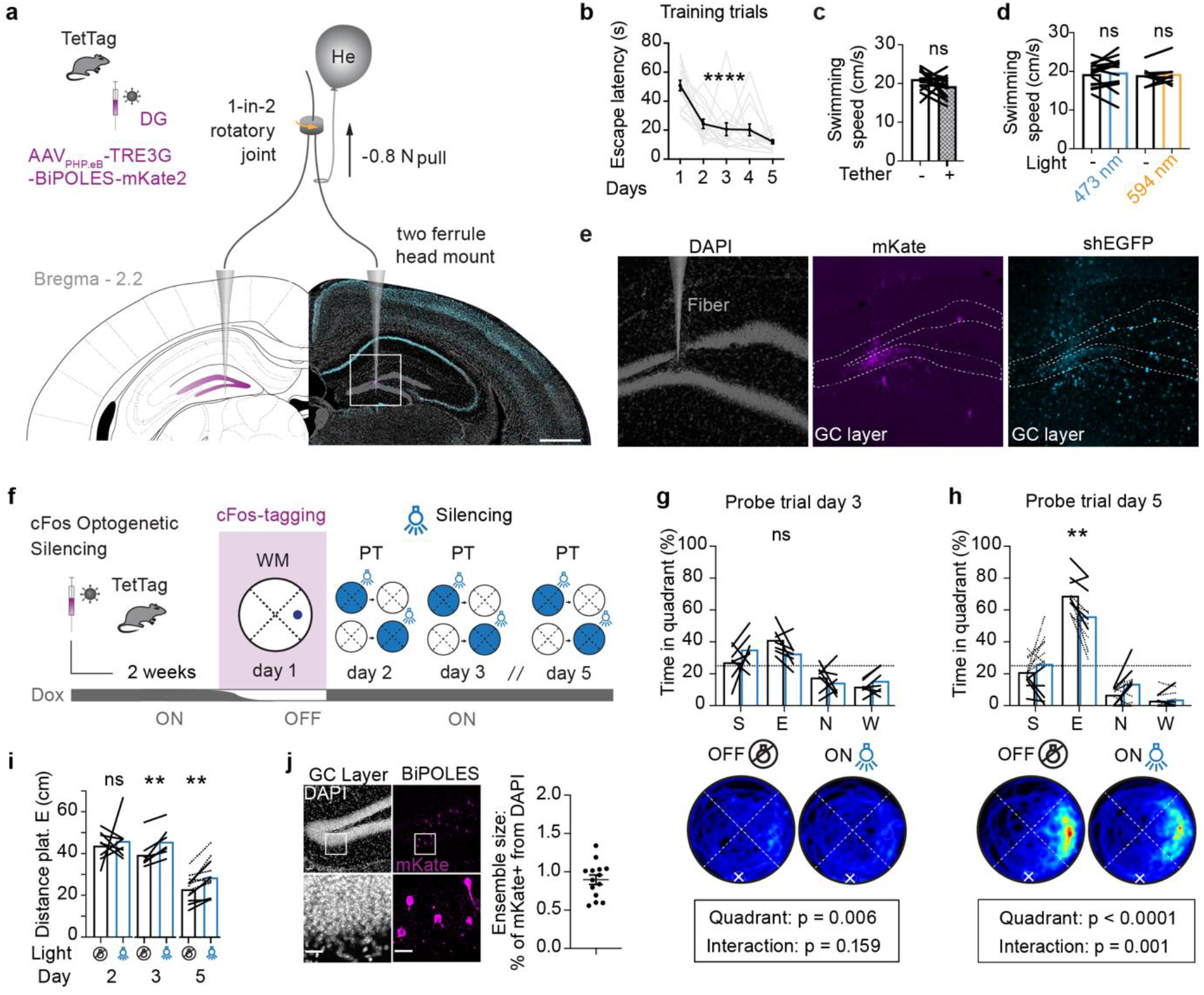
Optogenetic silencing of cFos^+^ granule cells impairs spatial memory recall. **a,** Tet-tag mice were injected with AAV for cFos-dependent expression of BiPOLES and red label (mKate2). Mice were bilaterally tethered with ultra-thin light fibers. **b,** Fibers and implant did not affect learning (black line: mouse mean, gray lines: individual mice, n = 15, ****, p<0.0001, one-way-ANOVA for repeated measures). **c**, Swimming speed of tethered mice with balloons was not different from the same animals in untethered trials (n = 15 mice, ns, p = 0.12, paired t-test). **d,** Optogenetic inhibition (473 nm) or excitation (596 nm) did not affect swimming speed (473 nm light, n = 15 mice, ns, p = 0.53, paired t-test) (594 nm light, n = 8 mice, ns, p = 0.74, paired t-test). **e**, Tapered fiber caused minimal damage in DG. **f,** On days 2, 3, and 5 of water maze training, the cFos-tagged ensemble from day 1 was inhibited in one of two tethered probe trials. **g,** On day 3, probe trial performance was poor, and optogenetic inhibition had no significant effect on the time spent in the target quadrant. **h,** On day 5, probe trial performance was very good, and optogenetic inhibition significantly decreased the time spent in the target quadrant (see Supplementary Table 1, two-way-ANOVA. Light x Quadrant interaction *** p = 0.001). Dotted lines are a second batch of mice. **i,** Optogenetic inhibition increased the average distance from the target platform location on days 3 and 5, but not on day 2 (see Supplementary table 1, paired t test: day 3 ** p = 0.004, Day 5 ** p = 0.002). **j,** After the behavioral experiments, expression of BiPOLES was verified in each animal (n = 15 mice).

We next tested whether silencing a larger percentage of GCs would impair spatial memory performance. Wild-type mice were injected bilaterally in the DG with AAV_9_-CaMKIIa-hM4Di-mCherry (non-conditional chemogenetic silencing group, n = 10). The hM4Di-mCherry expression in the DG strongly correlated with the effect of chemogenetic silencing on memory recall (R^2^= 0.67, p = 0.003, Supplementary Fig. 5). In strongly-expressing mice, chemogenetic silencing significantly impaired spatial memory recall. We estimate that silencing ~30% of GCs was required to decrease WM performance.

### Temporal dynamics of cFos ensembles in DG during spatial learning

We showed that the tagged ensemble from day 1 (cFos_d1_) must be activated when mice are performing in the WM, as optogenetic inhibition decreased performance on day 5. Does this mean cFos overlap continues to be high with the spacing of several days? Indeed, repeated cFos expression in DG has been reported in fear memory extinction ^39^; however, there is contradictory evidence when using Arc as a tagging IEG for memory extinction or retraining ^19,40^. We trained these mice with the same protocol that yielded the highest overlap from our previous experiments (RT group, Fig. 2a), but cFos-tagged the first ensemble on day 1 (Δt = 5 days) instead of day 5 (Δt = 1 day). In contrast to the overlap with an interval of 1 day, the overlap was not different from chance and significantly lower when Δt = 5 days (Fig. 4c). Thus, cFos ensembles are temporally unstable despite the stability of the environmental cues of the WM.

**Figure 4.**
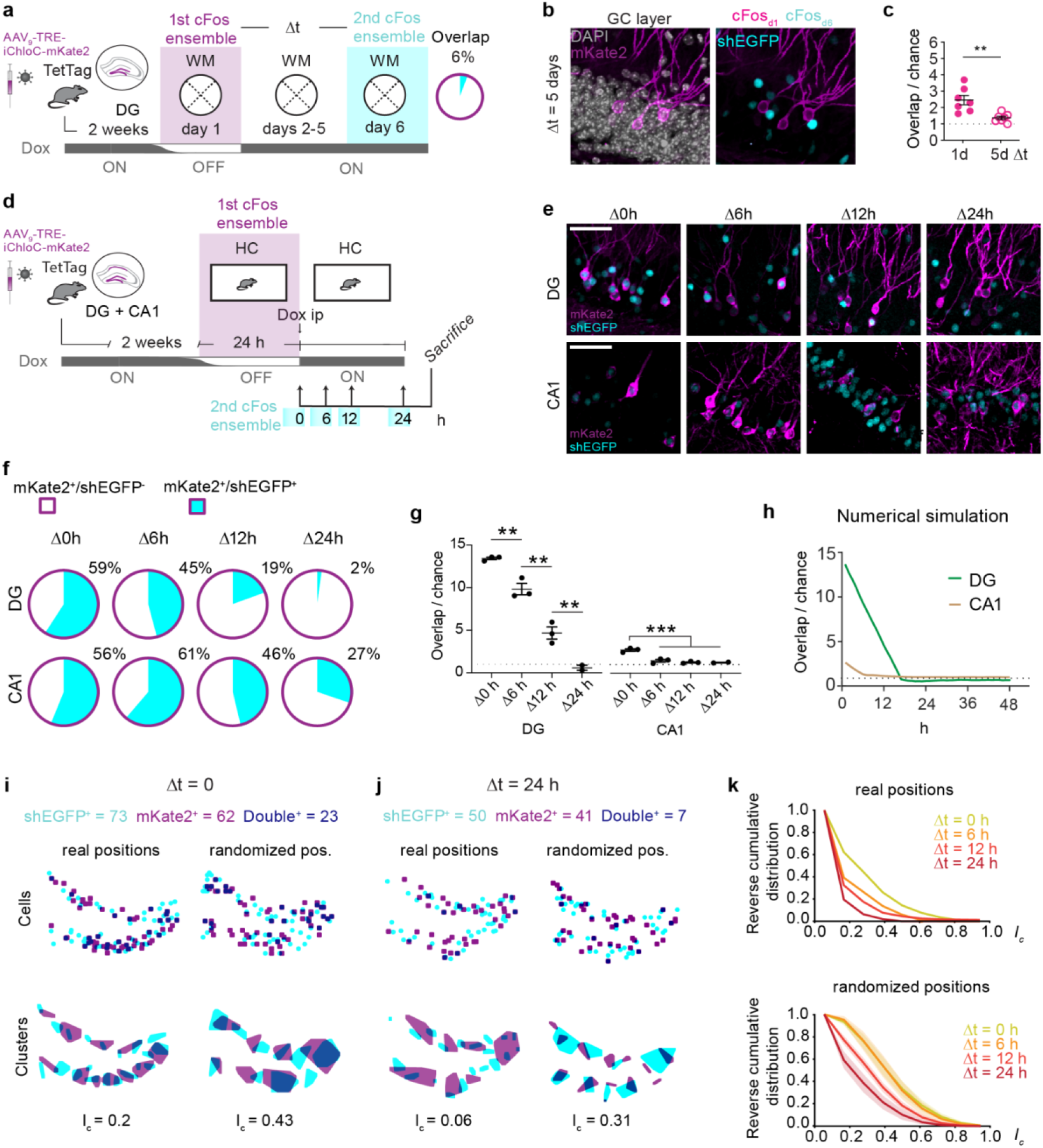
Temporal dynamics of cFos overlap in DG. **a**, TetTag mice were injected with AAV_9_-TRE-mKate2 (membrane) to tag cFos^+^ neurons from day 1 (cFos_d1_). They were trained for 6 days and sacrificed for cFos immunostaining. **b,** Immunofluorescence example images from the DG’s granule cell layer showing low overlap between cFos_d1_ (magenta) and cFos_d6_ (cyan). **c,** After 5 days of WM training, cFos overlap was not significantly different from chance (n = 7 mice, two-way-ANOVA, p = 0.11, see Supplementary Figure 2) and significantly lower (**, p = 0.003, unpaired t test) than after 1 day of WM training (RT group, data from Fig. 2d). **d**, Virus-injected Tet-tag mice were taken Dox-OFF in the home cage for 24 h and sacrificed at different time points after Dox injection. **e**, IF staining of cFos in DG and CA1. **f**, In DG, the fraction of the tagged ensemble (magenta) that was still positive for cFos (cyan) gradually decreased from 59% to 2% in the 24 h after closing of the tagging window. In CA1, the decrease was slower and less complete. **g**, Compared to the expected overlap for statistically independent groups (chance level), cFos patterns in DG started with extremely high overlap and dropped linearly to values below chance level. Pattern overlap in CA1 was only above chance when tested immediately at the end of the cFos tagging window (see supplementary table 1 for full statistical test. Δt effect: **** p<0.0001, two-way-ANOVA for both DG and CA1). **h**, Numerical simulation of cFos expression with negative feedback (brown curve) reproduces linear drop-in pattern similarity. CA1 simulation without negative feedback (green curve) does not drop below 1 (chance level)**. i,** Clusters of cFos^+^ cells have less overlap than expected by chance (randomized cell positions) at Δt = 0. **j**, At Δt = 24 h, cluster overlap has further decreased. **k**, Cluster overlap drops with increasing Δt. Clusters of real cellular positions (top panel) overlap less than expected by chance (randomized cell positions, bottom panel).

### In a stable environment, cFos ensemble selection is a function of time

In all our experimental groups so far, the overlap of cFos ensembles was relatively sparse (<10%). As the time difference (Δt) was always at least 24 h, we explored shorter intervals in a stable environment (home cage). Using a single injection at two different depths, we transduced both the DG and CA1 of TetTag mice with AAV_9_-TRE-mKate2. After two weeks, mice were OFF Dox for a 24 h period and sacrificed at 4 different time points: Immediately after the end of the OFF Dox period (Δ0h, n = 3), after 6 h (Δ6h, n = 3), after 12 h (Δ6h, n = 3) and after 24 h (Δ24h n = 3). All mice (except the Δ0h group) received an I.P. Dox injection to rapidly close the tagging window (Fig. 4d). Immediately at the end of the OFF Dox window (Δ0h), a high percentage of the mKate2^+^ cells were also positive for shEGFP (DG: 59%; CA1: 56%). During the next 24 h, this number gradually dropped to 2% in DG and to 27% in CA1, indicating different temporal dynamics of cFos expression in the two areas (Fig. 4e, f). We again calculated the overlap expected due to chance at the different time points and for CA1 (Supplementary Fig. 2, Fig. 4g). The results were consistent across animals, with DG dropping from 12-fold above chance to below chance level within 24 h (Fig. 4g). In CA1, the cFos ensembles were larger (Supplementary Fig. 2) increasing the expected overlap due to chance. Still the overlap was significantly higher than expected by chance in CA1 but only at the first time point (Fig. 4g). The very linear drop of pattern similarity in DG was initially surprising (Fig. 4g), but could be readily reproduced in a numerical simulation of a negative feedback loop, including the below-chance overlap at Δt = 24 h (Fig. 4h). The shEGFP reporter is more stable than cFos itself, suggesting that the pattern of endogenous cFos expression might change even faster. These results indicate that in a stable environment and without training (i.e., the home cage), a completely new set of cFos^+^ GCs is used for memory encoding on consecutive days.

### cFos-expressing GCs form spatial clusters that segregate over time

The sparse activity of GCs is controlled by robust inhibitory inputs from hilar interneurons ^41^. We were interested in the spatial patterns of cFos-expressing GCs, which may not be randomly distributed, but could instead reflect the sphere of influence of individual interneurons. We used the maps generated in the home cage experiments (Fig. 4d) to analyze clusters of cFos^+^ neurons at the different time points (Fig. 4i, j). As a measure of cluster overlap *I_c_*, we divided the overlapping area by the union of both areas (see methods). While at Δt = 0, clusters were partially overlapping (high *I_c_*), cluster overlap gradually decreased with time to very low values at Δt = 24 h (Fig. 4k). This result was not simply a consequence of the decreasing number of double-positive neurons because randomizing the positions of single- and double-positive neurons within the granule cell layer yielded clusters with higher overlap and less segregation over time. We conclude that the process of temporal segregation we describe here not only plays out in individual GCs, but also prevents re-use of DG regions that were highly active 24 h ago.

### GCs possess an intrinsic cFos blocking mechanism

The *in vivo* experiments revealed interesting differences between DG and CA1 concerning cFos dynamics. In principle, these differences could be due to temporal changes in synaptic input (i.e., network-level effects), or reflect differences in intrinsic properties, e.g. transcriptional or translational repression ^25,42^. We speculated that GCs are equipped with an intrinsic mechanism that prevents cFos expression on consecutive days. To test this idea, we used a chemogenetic approach to trigger cFos expression ^43,44^. When we transfected rat hippocampal slice cultures with a designer receptor coupled to Gq (DREADD, AAV_9_-CaMKIIa-h3MD(Gq)-mCherry), we found strong expression of cFos 70 min after clozapine-N-oxide (CNO) application (Fig 5a). A second CNO application 24 h later induced cFos only in a small number of GCs in the DG. In CA1 pyramidal cells, the second application was as effective as the first at inducing cFos expression (Fig. 5a). In addition to anti-cFos staining, we used a second antibody that recognizes FosB and its truncated splice variant ΔFosB. ΔFosB accumulates in neurons and can act as a suppressor of cFos in the DG ^25,42,45,46^. At the time of the second CNO stimulation, FosB/ΔFosB was strongly expressed in both DG and CA1 (Fig. 5b). The difference between the two hippocampal subfields was evident in the overlap analysis (cFos^+^/FosB^+^, Fig. 5c): Only 16% of DREADD-expressing GCs were cFos^+^ on both CNO stimulation days, compared to 59% of DREADD-expressing CA1 pyramidal cells, suggesting a cell-autonomous negative feedback loop that operates in GCs, but not in CA1 pyramidal cells. To check whether this mechanism is activated during WM training, we analyzed the brains of WM expert mice (Fig. 5d). On day 7, cFos expression was low in cells that in cFos-tagged neurons on day 1 (Fig. 5e, f). FosB/ΔFosB, on the other hand, could be clearly detected on day 7 in cells that had expressed cFos on day 1. Only 5% of the cells expressing cFos on day 7 were positive for FosB/ΔFosB, and only 3% of the high FosB/ΔFosB expressing cells had high cFos levels (Fig. 5g). Looking at individual mice, the overlap between cFos from day 1 and FosB from day 7 was 8 x higher than chance (Fig. 5h). ΔFosB protein is highly stable and known to accumulate in active neurons over time ^46,47^. Indeed, we found high and dense expression of ΔFosB in DG of overtrained mice (Fig. 5i). A long-lasting inhibition of cFos expression by FosB/ΔFosB could explain the decrease in cFos overlap over the course of training (Fig. 4c).

**Figure 5.**
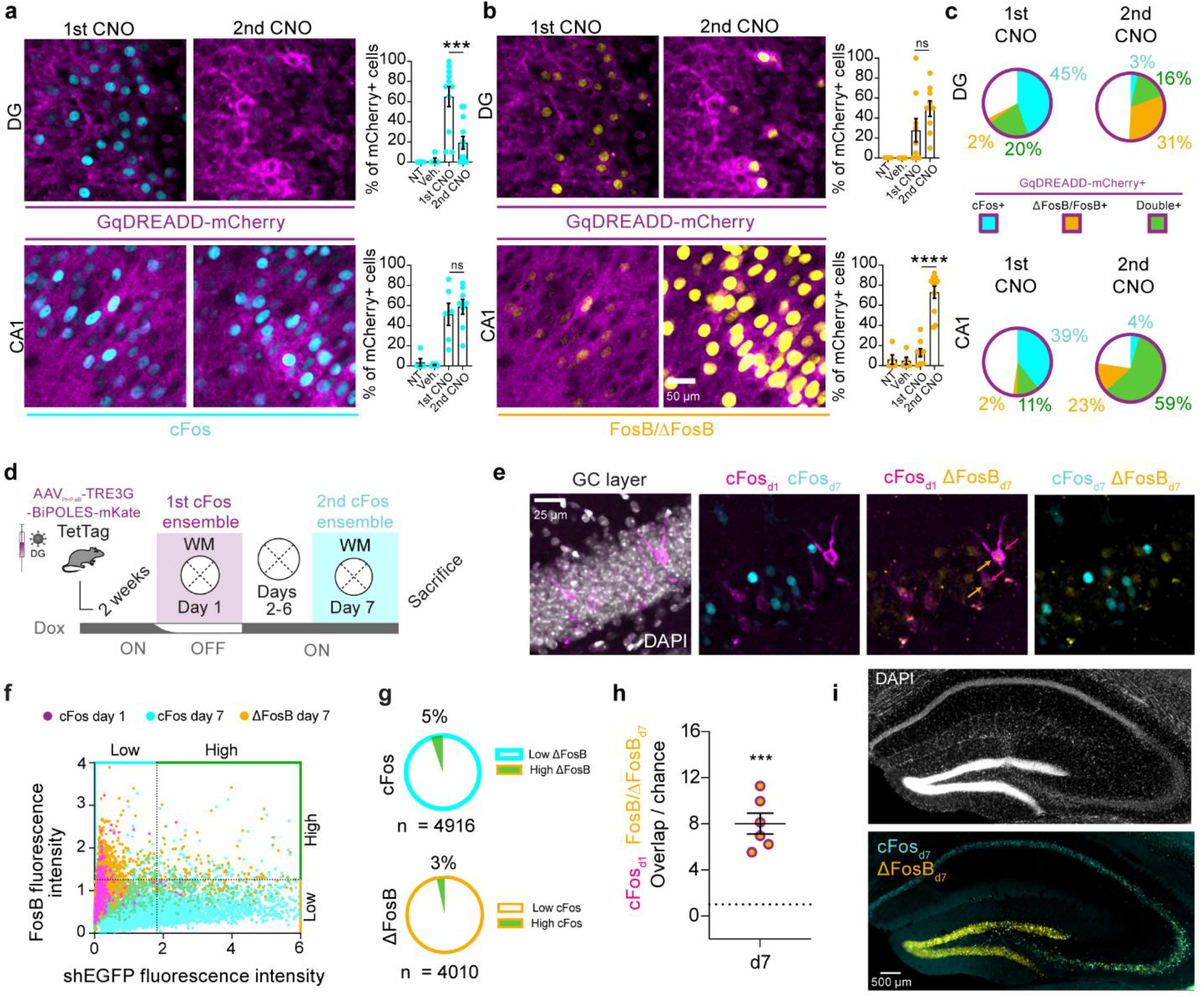
Mechanism of cFos suppression. **a** Chemogenetic activation of Gq in neurons to induce cFos expression in hippocampal slice culture. Gq-DREADD expressing neurons are identified by their mCherry fluorescence (magenta). Activation of Gq by CNO triggered cFos expression (cyan) on day 1 (1st CNO group). A second application of CNO on the next day (2nd CNO group) was much less efficient in triggering cFos expression in DG (upper row, 1st CNO vs. 2nd CNO group: ***, p = 0.0003), but equally efficient in CA1 (lower row, upper row, 1st CNO vs. 2nd CNO group: ns, p = 0.64). **b** Anti-FosB/ΔFosB staining (yellow) shows similar increased immunoreactivity on both days in DG (upper row, 1st CNO vs. 2nd CNO group: ns, p = 0.08). In contrast, CA1 shows only increased immunoreactivity on day 2 after the 2nd CNO stimulation (lower row, 1st CNO vs. 2nd CNO group: ****, p < 0.0001) (One-way-ANOVA. See Supplementary Table 1 for full statistics). **c** Overlap analysis of slice culture experiments. Of all DREADD-expressing cells (magenta), only 16% expressed both cFos and FosB (green) in DG after the second CNO application, but 59% in CA1. **d** cFos overlap analysis of day 1 vs. day 7 during the water maze training (Δt = 7 days. Mice from 1 batch of optogenetic manipulation experiments from Fig. 3, n = 6). **e** DG showing the cFos ensemble from day 1 (magenta), cFos ensemble from day 7 (cyan) and FosB/ΔFosB immunoreactivity (yellow). Note the double immunoreactivity of day1 cFos cells with FosB/ΔFosB (arrows), but no overlap between day 7 cFos cells and FosB/ΔFosB. **f** GC expression analysis pooled from 12 images per mouse. Little overlap between cFos expression on day 7 (x-Axis) and FosB/ΔFosB immunoreactivity (y-axis). cFos^+^ cells from day 1 (magenta) express FosB/ΔFosB, but not cFos on day 7. **g** Overlap analysis of data shown in f. Only 5% of day 7 cFos^+^ GCs show FosB/ΔFosB immunoreactivity. Only 3% of FosB/ΔFosB containing GCs were also cFos^+^ on day 7. **h** All mice displayed significantly high overlap than chance between day 1 cFos^+^ cells and FosB/ΔFosB on day 7 (*** p = 0.0004, paired t test). **i** Section of dorsal hippocampus from an expert mouse (6 days of WM training). DG shows strong accumulation of FosB/ΔFosB and sparse cFos expression, CA1 shows sparse FosB/ΔFosB expression and high cFos.

## Discussion

Fear conditioning experiments with optogenetic reactivation or suppression have created the notion that cFos-expressing neurons, a.k.a. engram cells, are the cellular representation of a memory as activating them is necessary and sufficient to create a state of fear ^17,48^. Compared to detecting the high-fear state as a lack of motion (freezing), spatial navigation is a more complex cognitive task that presents a richer read-out of goal-directed behaviors. In the WM, mice use visual information to estimate their current location relative to allothetic cues, which implies the flexible use of a cognitive map ^49,50^. For successful navigation, one’s own position and heading must be compared to the memorized location of the hidden target platform, a task that is highly dependent on intact hippocampal function ^2,51^. It takes a few days of training for a mouse to reach expert performance from an initially random search strategy (Fig. 1). In all phases of WM training, we found elevated levels of cFos in DG GCs (Fig. 2). The pattern of cFos^+^ GCs changed from day to day, even in expert mice. This is not what we expected, given the extremely high stability of place cells in DG ^10^. This raises two questions: What can cFos expression actually tell us about neuronal activity, and second, are cFos^+^ GCs place cells?

While cFos is known to be driven by high frequency spiking ^26,52^, our data suggest that the reverse is not true in the DG: the absence of cFos cannot be taken as evidence for the absence of activity. This became clear when we inhibited the activity of the original (day 1) cFos-tagged ensemble on consecutive days: WM performance was compromised in trials with optogenetic inhibition, even though only a very small fraction of GCs in dorsal DG (~ 1 %) was silenced (Fig. 3). The inhibition of cFos-tagged neurons was specific, as inhibition of random GCs was much less effective, we observed behavioral effects only in animals where > 30% of GCs were inhibited. Thus, GCs that expressed cFos on the first training day remained particularly relevant for successful navigation in that environment, even though they rarely re-expressed cFos on the successive training days. This finding poses a problem for the current definition of engram cells, stipulating “reactivation” in the same context as a defining criterion^17,19,39^. As the cFos-tagged neurons appear to be electrically reactivated in the absence of cFos “reactivation” (i.e. expression on subsequent days), using cFos as a proxy for electrical activity is problematic. Defining only the few GCs that express cFos twice as the “engram” cells will strongly underestimate the number of neurons relevant for memory recall. The DG cFos^+^ map produced on a given day will mostly contain GCs that have been activated for the first time in the WM. Thus, the DG cFos^+^ map may encode new features being added to the internal representation, possibly acting as a timestamp on the most recently acquired memory.

Why do reactivated GCs not express cFos? In slice cultures, DG GCs were refractory and did not readily express cFos 24 h after a first chemogenetic induction whereas CA1 neurons readily expressed cFos on successive days. FosB was also expressed after stimulation in both GCs and CA1 neurons, appearing quickly in DG and 24 h after stimulation in CA1. The FosB splice variant ΔFosB is known to be expressed in GCs and to inhibit expression of cFos in these neurons ^42^. Thus, ΔFosB may be the factor that renders DGs refractory to cFos on successive training days. Indeed, there was high overlap of FosB with the day 1 cFos-tagged GCs after 7 days of training (8x greater than chance), while FosB and cFos were almost mutually exclusive (Fig. 5). Both full-length FosB and the truncated variant ΔFosB are recognized by the antibody, but only ΔFosB is sufficiently stable to accumulate over days in neurons ^42,53,54^. It is likely that the variant that accumulated in the DG over multiple training days was primarily ΔFosB.

Why we observed no refractory period in CA1 of slice cultures is unclear: possibly, ΔFosB is not produced together with FosB in CA1 neurons, or an additional mechanism allows cFos expression to reoccur. We suspect that ΔFosB is not produced in CA1 pyramidal neurons as FosB staining did not appear to accumulate over days in these neurons in WM-expert mice. Indeed, in vivo imaging experiments have shown that in 60% of the cFos-expressing pyramidal neurons in CA1^43^ and 80% in barrel cortex^29^, cFos expression is not transient, but sustained for several days. We likewise observed that CA1 neurons express cFos rather persistently, with around 30% of the home cage cFos-tagged neurons continuing to express cFos 24 h after the mice were back ON Dox.

A new picture of time-controlled cFos regulation in the DG begins to emerge. In contrast to the CA1, only 19% of cFos-tagged GCs in the DG still express cFos after 12 h, and after 24 h, the overlap dropped below chance level. When neurons express cFos they are hyperexcitable ^28,55,56^, whereas overexpression of ΔFosB reduces excitability ^57^. Hyperexcitability increases the likelihood of burst firing and is ideal for inducing synaptic potentiation of incoming synapses ^58^. Burst firing also maximizes the impact of GCs on postsynaptic CA3 neurons, as the GC-CA3 mossy fiber synapses display extremely strong short-term facilitation ^59,60^. During the high cFos period, the probability of long-term potentiation may therefore be increased at both GC input and output synapses, i.e. onto CA3 pyramidal cells and interneurons targeted by the current set of cFos^+^ GCs ^61^. In contrast, synaptic plasticity may be reduced during the refractory period when ΔFosB is high, cFos expression is inhibited and excitability is reduced, possibly write-protecting the DG. A similar biphasic regulation of neuronal excitability in the amygdala has been linked to phosphorylation of CREB and is reversed by subsequent expression of the inhibitory CREB isoform ICER^62,63^.

Is it possible that the day 1 cFos-tagged ensemble is mainly composed of place cells? We would expect that the first time mice experience the WM, a set of place cells is created to tile this novel environment as shown in CA1 ^64^. These place cells should retain their place field over several days ^10^. While we were surprised overlap was not higher, there is above-chance cFos overlap in GCs of mice when re-exposed to the WM 24 h later, and no overlap in mice which instead explored a novel environment. Whether the few cFos-tagged GCs that expressed cFos the following day are place cells is unknown. Of note, when GCs are cFos-tagged on day 1, performance is quite poor and it is unlikely that a precise memory of the platform position had formed at this time. Our working hypothesis is that the mice formed WM-specific place cells on day 1, but not a specific spatial memory of the platform location. Performance was therefore degraded by inhibiting the WM-specific place cells, not by preventing recall of a specific spatial memory (i.e. the platform position). Further refinements of the WM task, e.g., by creating several escape options to choose from, could help to distinguish between these scenarios.

We propose that the assumption that cFos is an indicator of highly active ‘engram’ neurons may hold for naive mice housed under standardized (deprived) conditions and exposed to fear conditioning or other forms of one-trial learning ^65^. Multi-day training, which may be slightly closer to the daily challenges faced by mice in the wild, leads to the accumulation of ΔFosB in many GCs. Our results suggest that under these conditions, cFos patterns in DG depend not only on input from e.g. the entorhinal cortex, but very strongly on the activity history of each GC. Further investigation of memory systems “under load” during complex behaviors ^66^ may change our view of hippocampal processing and engram formation.

## Methods

### Experimental animals

B6.Cg-Tg^(Fos-tTA-Fos-EGFP*)1Mmay/J^ (TetTag) mice were obtained from the Jackson Laboratory (Strain #018306) and bred heterozygous crossed with C57BL6/J mice. Mice were group-housed with littermates until 2 weeks before rAAV injections, then were single-caged. Mice had access to food and water *ad libitum* and were kept in an animal facility next to the behavioral rooms on a reversed light-dark cycle (dark 7 am - 7 pm). All behavioral experiments were done during the dark phase of the cycle. Due to the weight of the implant and patch-cords (optogenetics), only male mice, between 20-40 weeks (>28g by the time of surgery) were included in the OptoWM experiments (Figure. 3–5). Both male and female mice were included in the cFos ensemble overlap experiments (Figure. 1, 2 & 6). All experiments were conducted in accordance with the German and European Community laws on protection of experimental animals and approved by the local authorities of the City of Hamburg (Behörde für Justiz und Verbraucherschutz, Lebensmittelsicherheit und Veterinärwesen, N 100/15 and N 046/2021).

### Viral constructs

To label the membranes of cFos^+^ neurons, an opsin ^67^ was cloned to a red fluorescent protein (iChloC-linker-mKate2). This construct was synthesized de novo and then inserted into a pAAV-TREtight backbone using MluI and EcoRI restriction enzymes to produce pAAV-TREtight-iChloC-mKate2. To drive spiking in cFos neurons, iChloC was replaced with CheRiff ^68^ to create pAAV-TREtight-CheRiff-mKate2. pAAV-TRE3G-BiPOLES-mKate2, a gift from Simon Wiegert, was used to inhibit cFos neurons. Constructs were packaged into AAV_9_ (iChloC-mKate2) or into AAV_PHP.eB_ (BiPOLES, CheRiff) by the UKE Vector Facility. AAVs for the DREADD experiments were ordered from Addgene (AAV_AAV_-CaMKIIa-h4MD(Gi)-mCherry, Addgene # 50476-AAV_9_-AAV_9_-CaMKIIa-hM3Dq-mCherry, Addgene # 50477-AAV_9_).

### Stereotactic injection and fiber implant

TetTag mice were virus-injected under analgesia and anesthesia using a stereotaxic drill and injection robot (Neurostar). Mice were fixed to the frame under isoflurane anesthesia (1.5% mixed in O_2_), skin and connective tissue was removed, and two craniotomies were performed using an automated drill on the desired coordinates. AAVs were injected at 10^12^ vg/ml concentration, except for iChloC-mKate2 experiments where it was 10^13^ vg/ml. AAVs were delivered bilaterally into the dorsal hippocampus using a glass micropipette attached to a 5 μl syringe (Hamilton). A single injection per site was performed using stereotaxic coordinates for DG (−2.2 AP, ± 1.37 ML, −1.9 DV) with a volume of 500 nl on each side (injection speed: 100 nl/min). CA1 was also injected in a set of experiments (Fig. 5) by moving to −1.4 DV and injecting 400 nl after the DG injection. After the last injection, the pipette was retracted 200 μm and left for at least 5 min to minimize retroflux. After the injections, the bone surface was cleaned with 0.9% NaCl solution and the skin was stitched. To avoid hypothermia, a heated pad was placed under the animal during surgery and under its cage for 1 h until full recovery. We provided post-surgery analgesia with Meloxicam mixed with softened Dox-food (see below) for 3 days after surgery. Animals recovered at least 2 weeks before behavioral experiments. Mice for optogenetic experiments were implanted with a custom-made bilateral tapered tip optic fiber implant (Doric) targeting the DG sulcus (−2.2 AP, ± 1.37 ML, −1.7 DV) right after AAV injection. Implants were attached using dental cement (C&B Metabond) onto the skull’s surface and a protective cap was made out of an Eppendorf tube. The cap was secured then by applying acrylic resin (Pattern Resin LS, GC America) on all the exposed skull’s surface in the implant surroundings.

### Doxycycline treatment

Animals were given doxycycline-containing food (Altromin-Dox, 50 mg per kg of body weight, red pellets). To tag the first cFos ensemble, animals were changed to doxycycline-free food (Altromin, light-brown pellets) 24 h before exposure to the task. To ensure that the animals did not eat any Dox-food crumbles that had fallen into their cages, they were moved into new cages with fresh bedding when food was changed. The old nesting material was transferred to the new cage to decrease novelty. Dox-food was resupplied exactly 24 h after removal, right before the behavioral task. For cFos ensemble temporal shift experiments (Fig. 4), an I.P. injection of doxycycline (50 μg / g body weight) was injected after the end of the 24h OFF Dox period.

### Behavioral experiments

For ensemble overlap experiments, mice were injected in batches and randomized to each of the groups. Optogenetic experiments had a crossover design where all mice had probe trials with and without light, to minimize sampling errors and reduce the number of animals required to reach statistical power (paired statistics). Mice that did not have adequate viral transduction (see image analysis section), had off-target implants or performed poorly (floaters or implant intolerance) were excluded from the analysis. All experiments were recorded on digital video and automated mouse tracking was done with Ethovision XT 11.5.

#### Water Maze (WM)

Animals were handled for 1 week before the start of the pre-training sessions to reduce stress during behavioral tasks. **Pre-training.** Mice were pre-trained for 2 days before their first exposure to the WM arena. Sessions (3-4 trials of max. 60 s each on two days) were done in a small rectangular water tank in the dark, in the same room where the WM task was performed. Water level was 1 cm above the 14 cm diameter escape platform. The position of the platform was alternated between the left and right side of the tank between trials, keeping a distance of 5 cm from the walls to avoid thigmotaxis. Once the animals found the platform, a grid was presented until the animals climbed onto it and were returned to their home cages in the waiting area of the behavioral room. **Training.** The WM consisted of a circular tank (1.45 m diameter) with visual asymmetrical landmarks, filled with water mixed with non-toxic white paint to prevent the animals from seeing the platform (submerged by 1 cm). The platform was placed in the center of the east quadrant during regular training and switched to the opposite (west) quadrant for reversal training (max. swim time: 90 s). To test spatial reference memory, a probe trial without a platform was performed on each day. Mice underwent six trials every day (4 training trials (TT, 90 s) + 1 probe trial (PT, 60 s) + 1 TT), inter-trial interval (ITI) was 8-10 s. For the TTs, mice were lowered into the tank facing the wall in different, pseudo-randomized positions (avoiding the target quadrant). In the PT, mice were lowered in the center of the tank. An opaque cup-sized chamber attached to a pole was used to transfer the mice from their home cage to the drop position and a plastic grid attached to a pole was used to pick up the mice. Mice were picked up 10 s after they found the platform and were returned to their home cage. Mice that did not find the platform during the TT were guided to it using the grid and were picked up after a 10 s on-platform waiting period. For both pre-training and WM, water temperature was 19-21°C and a heat lamp was placed over the waiting area to avoid hypothermia.

#### Home cage and open field control experiments

To evaluate the temporal effect of cFos ensembles, mice were unperturbed in their home cage (HC) during the OFF Dox period, but no behavior was analyzed. Mice were either sacrificed right after 24h from Dox removal (Δ0 group) or put back on Dox and sacrificed after 6, 12 or 24 h. For all the cFos overlap experiments, both cFos^+^ tagging events were designed to have the same amount of trials and training length. However, for the open field experiment, mice were placed inside a square arena (50 x 50 cm, 50 lux) for 20 min but were kept in the behavioral room the same time as their WM counterparts.

#### Optogenetic silencing in the water maze

Mice were connected to two thin optic fibers (200 μm core diameter, 2 m, Doric Lenses, Canada) using ceramic ferrules (Doric) and put back to their home cage for at least 5 min before any behavioral task. In pre-training trials (2 days, 3 trials per day), mice were connected and disconnected before every trial to habituate them to the tether. To avoid stress-related cFos tagging, mice were never tethered during the Dox-OFF period. In all trials with tether, the weight of the fibers was compensated with a white helium balloon (0.08 N pull force), attached to a light fiber ~30 cm above the mouse with a transparent thin plastic tubing, long enough to keep the balloon out of the field of view of the video camera. Optogenetic silencing occurred only during memory recall (no target platform) probe trials (PTs). Blue light was delivered using a laser combiner (LightHUB, Omicron) connected to a commutator (FRJ_1×2i_FC-2FC_0.22 Doric Lenses, Canada) to split the output into two fibers. The commutator was placed close to the ceiling, at the same height as the camera. To ensure inhibition, blue illumination was started when mice were still in their HC (473 nm light pulsed at 33 Hz, 10 ms pulses, 8-10 mW). Immediately after light *on*, mice were placed on a starting platform facing the wall on the south quadrant. The starting platform was submerged with a hydraulic mechanism (Atlantis platform) after 30 s. After 60 s of swim time, the target platform (east quadrant) was raised to just below the water level. Mice that found the target platform were transferred to their HC 10 s later. In cases where mice did not find the platform they were guided to it and then transferred to their HC. Blue light was *on* for a total of 2 min, covering the entire time inside the WM tank. Probe trials (no light) were done exactly the same, but with the blue laser *off*.

#### Training protocol

Two batches of mice were used for optogenetic manipulation in the water maze. On training day 1, the protocol consisted of six trials: four training trials of max. 90 s with the platform in quadrant E, followed by a probe trial (60 s, without platform) and another training trial. On day 2, mice were placed on the starting Atlantis position for 90 s (visual recall trial). Afterwards they performed two probe trials while blue inhibiting light was being delivered in the same cross-over design, followed by three training trials. Day 3 started with two training trials, followed by two probe trials (cross-over design, 473 nm light pulses), and two more training trials. On day 4 mice underwent the standard protocol (four training trials, one probe trial without light, one training trial). Day 5 training, mice had two training trials followed by two probe trials (473 nm light, cross-over design). We repeated this protocol on day 6, replacing 473 nm inhibiting light pulses with 594 nm exciting pulses during probe trials. This batch was perfused for brain slicing and immunohistochemistry on day 9. The second batch was trained similarly. After tagging (training day 1) mice went through the same training protocol from day 2 to day 4, only adding a reactivation trial before the first training trial. During this reactivation trial, mice were connected to the laser via an optical fiber on the starting platform, delivering 594 nm light pulses (or no light for control mice) for 90 s. Optogenetic stimulation had no effect on behavior, nor cFos overlap, nor FosB expression so we pulled the data from both light and no light groups together. On day 5, these mice had two training trials, followed by two probe trials, where half of the mice received 473 nm light pulses for the duration of the trial. On day 6 mice had two more training trials and a probe trial. This batch of mice was perfused for brain slicing and immunohistochemistry on day 7, exactly 90 minutes after performing in one probe trial without light.

#### Chemogenetic silencing in the WM

Wild-type mice were bilaterally injected in DG with AAV_9_-h4MD(Gi)-mCherry. After at least 2 weeks of recovery, mice were handled and pre-trained like the previous groups. On day 1, mice were trained for 6 trials. On day 2, mice had two probe trials (PT). Forty minutes before the 1st PT, they received an intraperitoneal (I.P.) injection of 0.9% NaCl solution (Vehicle). Mice were then injected with clozapine-N-oxide (CNO, 5mg kg^-1^) diluted in 0.9% NaCl solution and forty minutes later, they did the 2nd PT. From day 3-7 mice were trained every day with 4-6 TT and 1 PT. On day 8, mice were injected again with CNO forty minutes before a PT. A subset of mice was sacrificed to confirm the silencing effect of CNO. The rest of the mice were sacrificed on subsequent days for immunohistochemistry.

### Ex vivo brain processing and cFos immunofluorescence (IF) staining

Mice from all cFos overlap analysis experiments were perfused 3 h after the last trial and mice in the optogenetic silencing were perfused 1.5 h after the last trial. Mice were injected with ketamine/xylazine (100/10 mg kg^-1^) intraperitoneal and intracardially perfused with 1x phosphate-buffered saline (PBS, Sigma) followed by 4% paraformaldehyde (PFA, Roth). Brains were extracted and stored in 4% PFA for at least 24 h at 4°C. Before sectioning, brains were washed with 1xPBS for 20 min at room temperature (RT). The dorsal hippocampal region (−1.2 to −2.3 AP from bregma) was cut in 40-50 μm coronal sections using a vibratome (Leica VT100S) and collected in PBS. From each series, six sections were selected from Bregma −1.7 to −2.3 mm and incubated in blocking buffer (1x PBS, 0,3% TritonX, 5% goat serum) for 2 h at RT. Next, sections were placed in the primary antibody carrier solution (1x PBS, 0,3% TritonX, 1% goat serum, 1% BSA) and incubated overnight at 4°C. After 3x washing with 1xPBS for 5 min, the sections were incubated for 2 h at RT in the secondary antibody carrier solution (1x PBS, 0,3% TritonX, 5% goat serum). Sections were washed 3 x 10 min in 1x PBS, stained with 4’,6-diamidino-2-phenylindole (DAPI, 1:1000) for 5 min and mounted on coverslips using Immu-Mount (Shandon). Primary antibodies: Chicken Anti GFP polyclonal antibody, Invitrogen (A10262, Lot 1972783); Rabbit Anti-tRFP (mKate2) antibody, Evrogen (AB233, Lot 23301040466), Rabbit anti cFos, Synaptic Systems (226003), Rat anti cFos, Synaptic Systems (226 017, Lot 1-6), Rabbit anti FosB (5G4), Cell Signaling Technology (#2251, Lot 3). Secondary antibodies: Alexa Fluor 488 conjugated secondary antibody (goat anti-chicken, 1:1000, Life technologies; A11039; Alexa Fluor 568 conjugated secondary antibody (goat anti-rabbit, 1:1000, Life Technologies, A11011).

### Confocal imaging

Before imaging, mounted slides were labeled with a code to blind the analyst. For cFos overlap experiments, the stained slices were imaged with a confocal microscope (Olympus Fluoview FV 1000) using an oil immersion objective (UPLSAPO 20X NA: 0.85). 15μm were imaged from each of the 6 sections per brain (10 image stack at 1024 x 1024 resolution, Z-step: 1.5 μm) was acquired from the left and right dentate gyrus. For all the experiments done with BiPOLES. slices were imaged with a Zeiss Confocal microscope LSM 900 using an air objective (APOCHROMAT, 20X, NA: 0.8). For these images 18 μm were imaged (6 image stack at 1024 x 1024 resolution, Z-step: 3 μm). Excitation/emission filters were selected using the dye selection function of the Fluoview/Zen3.5 software (Alexa 405 (DAPI), Alexa 488 (shEGFP) and Alexa 568 (mKate2), Alexa 647 (cFos or FosB). The image acquisition settings were optimized once and kept constant for all images within an experimental data set. Images were not deconvolved, gamma-adjusted or filtered. Example images presented in figures were median-filtered (2×2 kernel) and cropped. Each color channel was linearly adjusted (Image J).

### Ensemble size calculation and overlap analysis

Confocal image stacks (pseudo-colored cyan: shEGFP; pseudo-colored magenta: mKate2; pseudo-colored gray: DAPI) were analyzed using Imaris (Bitplane). The DAPI channel was used to create a surface of the upper and lower blade of DG (granule cell layer, GCL). The volume of the GCL was used to estimate the total GC number based on published cell densities^49^ and to mask the cyan and the magenta channel to restrict analysis to the GCL. Automatic spot detection was used to identify shEGFP^+^ cells in the cyan channel. The quality filter (round nuclei with a diameter of 8 μm) was adjusted once and then applied to every image stack of the same experiment. False positive spots (e.g. staining artifacts) were manually removed. It was not possible to detect mKate2-expressing GCs automatically as only the plasma membrane was labeled. To count mKate2-positive cells, spots were placed manually (spot size 12 μm) using the pseudo-colored magenta channel only. Double-positive cells (distance between shEGFP^+^ and mKate2^+^ spots < 5 μm) were identified using a Matlab script. They were then manually inspected to check for artifacts and, if necessary, corrected. From the Imaris analysis, we calculated the following quantities:

1. number of qranule cells 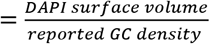
2. fraction of shEGFP taqqed cells 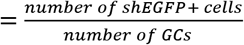
3. fraction of mKate2 taqqed cells 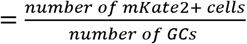
4. fraction of double positive cells 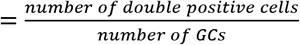
5. expected overlap = fraction of shEGFP cells * fraction of mKate2 cells
6. Overlap / chance 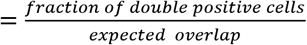

### Cluster analysis

Neurons were clustered by the Agglomerative Clustering algorithm implemented in the scikit-learn python library. For each image, we first computed the connection graph by using kneighbors_graph, which is also implemented in scikit-learn. The number of neighbors was fixed as one-tenth of the total number of neurons. First (mKate^+^) and second set (shEGFP^+^) of cFos-positive neurons were clustered separately (linkage criterion: “ward”/ Clustering was performed for 16 different numbers of clusters *n_c_* ∈ [5,20]. To determine the optimal number of clusters (*n_c_*) for each image, we generated 500 sets of artificial data from each image by randomly distributing the cFos^+^ neurons within the cell body layer of DG. Double-positive neurons were treated as a third type, keeping their numbers identical to the experimental data. The cell body layer area was defined by the DAPI channel, and the size of each neuron was standardized (16*16 pixels). Each set of artificial data was clustered with the same procedure as the empirical data. For each clustering result, a clustering index

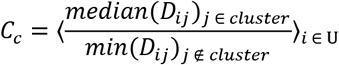

was calculated, where *D_ij_* is the distance between neurons *i* and *j*. Smaller *c_c_* indicates stronger clustering. For each *n_c_*, we calculated the proportion (of the 500 artificial sets) where *c_c_* was smaller in the artificial data than in the empirical data. The optimal *n_c_* was defined as the one with the smallest proportion, i.e. the strongest clustering compared to randomly distributed cells. In the case of multiple *n_c_* with the same smallest proportion, the largest *n_c_* was selected. The optimal *n_c_* was determined for mKate2^+^ and shEGFP^+^ clusters separately (always including the double-positive cells).

The sphere of influence of a cluster was defined as the convex hull of all its neurons, calculated by using ConvexHull implemented in SciPy python library. The overlap index between mKate2^+^ clusters *M_i_* and shEGFP^+^ clusters *S_j_* was defined as *I_c_* = (*M_i_* ∩ *S_j_*)/(*M_i_* ∪ *S_j_*). In case of overlap between multiple clusters, the largest overlap was scored. Another 200 sets of artificial data were generated from each image and analyzed in an identical fashion to create cumulative distributions of *I_c_* values expected for randomly distributed cFos^+^ cells (Fig. 5h).

### Hippocampal slice cultures

TetTag mice or Wistar Unilever rats (Envigo, HsdCpb:WU) were prepared at postnatal day 4–8 as described ^69^. Briefly, animals were anesthetized with 80% CO_2_ 20% O_2_ and decapitated. Hippocampi were dissected in cold dissection medium containing (in mM): 248 sucrose, 26 NaHCO_3_, 10 glucose, 4 KCl, 5 MgCl_2_, 1 CaCl_2_, 2 kynurenic acid, 0.001% phenol red (310–320 mOsm kg^-1^, saturated with 95% O_2_, 5% CO_2_, pH 7.4). Tissue was cut into 400 μM thick sections on a tissue chopper and cultured on membranes (Millipore PICMORG50) at 37 °C in 5% CO_2_. No antibiotics were added to the slice culture medium which was partially exchanged (60–70%) twice per week and contained (for 500 ml): 394 ml Minimal Essential Medium (Sigma M7278), 100 ml heat inactivated donor horse serum (H1138 Sigma), 1 mM L-glutamine (Gibco 25030-024), 0.01 mg ml^-1^ insulin (Sigma I6634), 1.45 ml 5 M NaCl (S5150 Sigma)), 2mM MgSO4 (Fluka 63126), 1.44 mM CaCl_2_ (Fluka 21114), 0.00125% ascorbic acid (Fluka 11140), 13 mM D-glucose (Fluka 49152).

### High K^+^ stimulation

To characterize cFos (native) and shEGFP (transgene) expression, TetTag mouse organotypic slice cultures were stimulated with a 50 mM KCl solution as described ^52^. In brief, slices were submerged in high KCl solution for 2 min, 3x with an interval of 10 min, alternating with HEPES (see below). Slices were washed in HEPES solution before returning to the incubator. Slices were then fixed for 30 min in 4 % PFA starting at 0, 30, 60, 120, 240 mins after stimulation subsequently processed for cFos and shEGFP immunofluorescence staining as above. After staining, slices were imaged in a confocal microscope and immunoreactive cells were counted using an automated spot detection (Imaris) tool (see above).

### cFos expression induced by chemogenetic stimulation

Organotypic rat slice cultures were injected with AAV_9_-CaMKIIa-hM3Dq-mCherry at DIV 14 using a Picospritzer III (Parker Hannafin). The virus was injected in DG, CA3 and CA1. The pressure was set to 1.8 bar with a pulse duration of 50ms. After 4-5 days of expression, slices were treated either once or twice with CNO (1 μM, TOCRIS) 24 h hours apart. Slices were stimulated by applying a 10 μL drop of CNO diluted in HEPES (in mM: 125 NaCl, 25 NaHCO_3_, 25 C_6_H_12_O_6_, 1.25 NaH_2_PO_4_, 2.5 KCL, 1 MgCl, 2 CaCl_2_, pH 7.4) on top (1st CNO group). Some of these slices were then fixed in 4 % PFA. Another set of slices was transferred to a new well containing only HEPES for 3 minutes to wash off the CNO and then returned to the incubator. These slices were then stimulated again the following day by a drop of CNO solution (2nd CNO group). This time, diluted in HEPES plus TTX (1-10 μM) to ensure a cell autonomous activity. As controls, some slices were never treated (NT group) or only treated with HEPES + TTX without CNO (vehicle group). All slices were fixed 70 min after the solution drop was applied and stained against cFos and FosB like described in the immunohistochemistry section. A z-stack was taken for each hippocampal subregion but analysis was performed in a single plane image at 20-25 μm from the slice surface. A manual (blinded) spot count was done using Imaris software. By visualizing only the mCherry channel, spots were manually set (20 spots for DG, 50 spots for CA1 per slice) in the nuclear region of pseudo randomly selected mCherry^+^ neurons. The cFos and FosB intensity data of all spots were exported and the analysis was done based on the intensity profile of the other two channels. All the spots were then sorted using a Matlab script into cFos^+^, FosB^+^, double^+^ cells. The Intensity threshold for cFos^+^ and FosB^+^cells was set manually and kept the same for all conditions of the same experiment.

### Acute slice preparation

Mice were decapitated under CO_2_ anesthesia and the brains dissected. Acute coronal slices (300 μm) were cut on a vibratome (Leica VT1000 S) in cold (4°C) cutting solution containing (in mM): choline-chloride (110), KCl (2.5), NaH2PO4 (1.25), NaHCO3 (25), MgCl2 (7), CaCl2 (0.5), glucose (25), sodium-ascorbate (11.6), sodium-pyruvate (3.1). The osmolarity of the choline solution was typically 310 mOsm/l, with a pH of 7.4. Oxygenation and pH were maintained by bubbling (95% O_2_, 5% CO_2_). Slices were kept in a holding chamber with artificial cerebrospinal fluid (aCSF) containing (in mM): NaCl (125), KCl (2.5), NaH_2_PO_4_ (1.25), NaHCO_3_ (26), MgCl_2_ (1), CaCl_2_ (2), glucose (10), saturated with 95% O_2_, 5% CO_2_. Brain slices recovered at 34 °C for at least 45 minutes before the start of experiments.

### Electrophysiology and optogenetic stimulation

Acute slices were superfused with aCSF at a rate of 2.5–3 ml / min at 30–31°C. Patch pipettes were pulled from borosilicate glass and had a resistance of 3–5 MΩ when filled with internal solution containing (in mM): K-gluconate (135), HEPES (10), MgCl_2_ (4), Na_2_-ATP (4), Na-GTP (0.4), Na_2_-phosphocreatine (10), L-ascorbic acid (3), EGTA (0.2). Internal solution had pH 7.2 and 295 mOsm/L. The mKate2-positive cells in the DG were patched and signals acquired through an Axopatch 200B or Multiclamp 700B (Axon Instruments, Inc.), National Instruments A/D boards and Matlab running Ephus software ^70^. Action potential firing was electrically evoked by somatic current injection. Optogenetic stimulation was given through the objective (Olympus, 60x, 1.0 NA) at 473 or 594 nm (CoolLED, pE-4000). Data were analyzed with Matlab or Clampfit 10.7 (Molecular Devices).

### Statistical analysis

The definition and the exact value for n is given in the text or in figure legends. Statistical analysis was performed using GraphPad Prism (v8). Performed statistical tests are described in each figure legend and more detailed in Supplementary Table 1. All tests were two-tailed and the level of significance was set at p < 0.05.

## Acknowledgments

We would like to thank Iris Ohmert, Jan Schröder, Katryn Sauter, Sabine Graf, Chantal-Joy Agbo, Tanja Stößner and Karen Kesseler for excellent technical assistance. The pAAV-TRE3G-BiPOLES-mKate2 construct was a gift from J. Simon Wiegert, Ingke Braren from the UKE Vector Facility produced rAAV. We thank Mary Muhia, Irm Hermans-Borgmeyer, Silvia Rodriguez-Rozada and Brenna Fearey for discussions and helpful advice.

## Author contributions

PLM, AF, AKE, FM and TGO conceived and designed experiments. PLM, AF, LB, and LA performed behavioral experiments. AF, TF, JA, LL and LB performed electrophysiological experiments. DL and CCH applied cluster analysis. AF and CEG performed slice culture experiments. PLM, LB, LA and FM analyzed behavioral data. Confocal imaging, immunohistochemistry and cFos scoring was done by AF and LB. PLM, MK, AKE, CCH, FM and TGO obtained funding and provided infrastructure. PLM, CEG, FM and TGO wrote the paper. All authors edited and approved the manuscript.

