## Supplementary Figures 1-6 for "cFos ensembles in the dentate gyrus rapidly segregate over time and do not form a stable map of space"

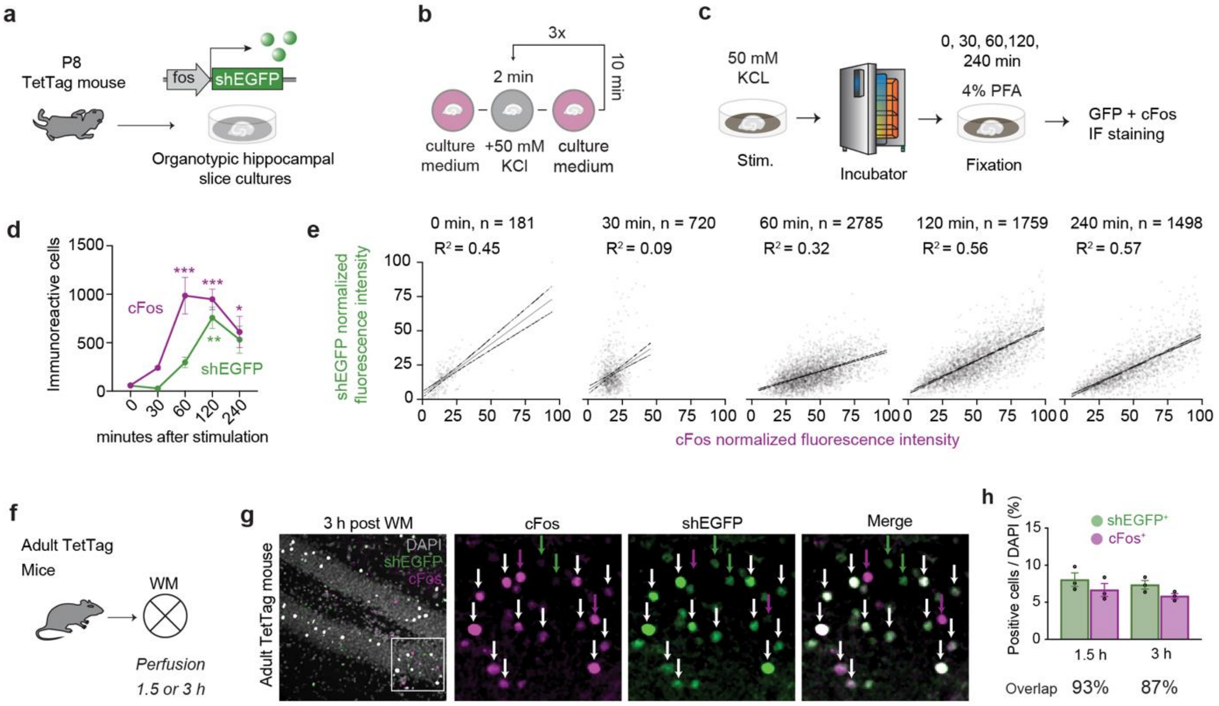


**Supplementary Figure 1. Characterization of cFos reporter expression in TetTag mice. a**, Hippocampal slice cultures were prepared from p8 TetTag mice in which shEGFP is expressed under the cFos promoter. **b**, cFos expression was induced by 50 mM potassium chloride (KCl) application. Slices were exposed to high KCl for 2 min, 3 times at 10 min intervals. **c**, Slices were returned to the incubator (37 °C) after stimulation and fixed with 4% paraformaldehyde (PFA) 0, 30, 60, 90, 120 or 240 min after the end of the stimulation. **d**, Time course of native cFos protein and shEGFP reporter expression. Immunoreactive (IR) cells were manually scored (shEGFP, n = 3 slices; cFos, n = 3 slices, mean ± SEM) inside the granule cell layer as determined by the DAPI signal. $cFos^+$ cells significantly increased after 60 min while $shEGFP^+$ cells significantly increased after 120 min (two-way-ANOVA, Time Point x Protein interaction * p=0.048) **e**, Anti-cFos vs. anti-shEGFP fluorescence intensity, using automatic detection of cFos+ cells (low detection threshold). High linear correlation was found 120 min and 240 min after stimulation. **f,** TetTag mice were trained in the WM to induce cFos and sacrificed 1.5 h or 3 h after the end of the last training session. **g,** Dentate gyrus from TetTag mouse stained against cFos (magenta) and shEGFP (green) 3 h after water maze (WM) training. Most fluorescent granule cells (GCs) are double-positive (white arrows). Magenta and green arrows correspond to $cFos^+$ or $shEGFP^+$ GCs, respectively. **h**, Fraction of positive cells and overlap of shEGFP reporter (green) and cFos protein (magenta) in animals sacrificed 1.5 h (n = 3) and 3 h (n = 3) after water maze training. Markers correspond to individual animals, bars show mean ± SE.

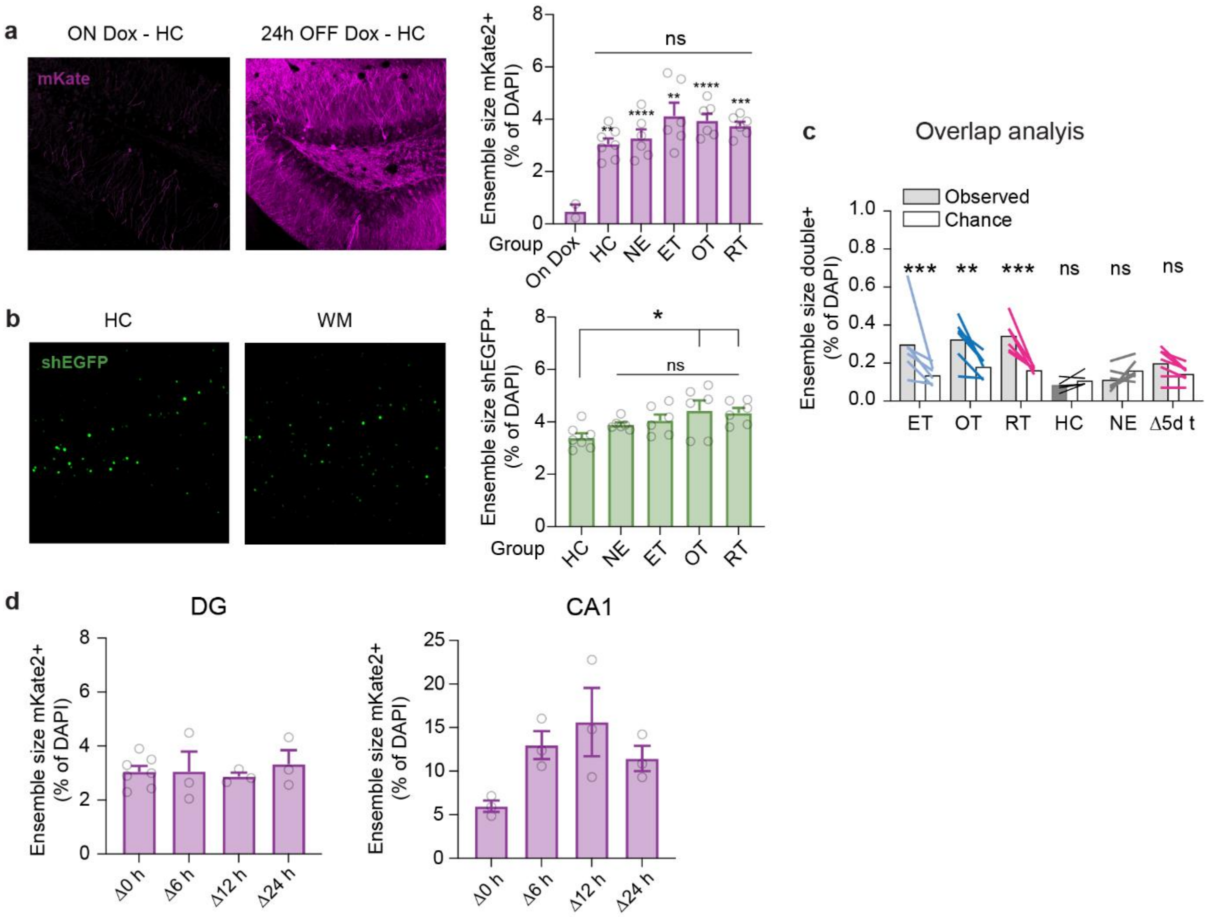


**Supplementary Figure 2. Size of cFos$^+$ ensembles in TetTag mice. a**, Removal of Doxycline food induced cFos-dependent expression of membrane-targeted mKate2 in the home cage (HC). The fraction of mKate2-positive neurons was not different in home-caged animals compared to novel environment (NE) and mice trained in the water maze (early training ET; overtrained OT; reversal training RT). All groups showed significantly larger cFos$^+$ ensemble sizes than mice on Dox (HC **, n = 7; NE ****, n = 6; ET ***, n = 6; OT ****, n = 6; RT ***, n = 6. One-way-ANOVA, Sidak multiple comparison test). **b**, Nuclear shEGFP reports cFos-expression in the 2-3 hours before sacrifice. Ensemble size was significantly larger in the OT and RT groups compared to the HC group. Ensemble size was similar in all WM-trained groups (HC vs. OT *, n = 6; HC vs. RT *, n = 6; HC vs. NE ns, n = 6; HC vs. ET ns, n = 6. One-way-ANOVA, Sidak multiple comparison test). **c**, Observed and chance-level overlap between mKate2 and shEGFP-tagged ensembles from different (Figs. 1, 2 and 4) experimental groups (HC ns, n = 7 (two sets of lines are overlapping); NE ns, n = 6; ET ***, n = 6; OT **, n = 6; RT ***, n = 6; Δ5d ns. Two-way-ANOVA, Sidak's multiple comparison test). **d**, Ensemble size (mKate2+) in home caged mice sacrificed at different times after the end of a 24 h off-Dox period (see Fig. 4). Note delayed increase of mKate2+ pyramidal cells in CA1.

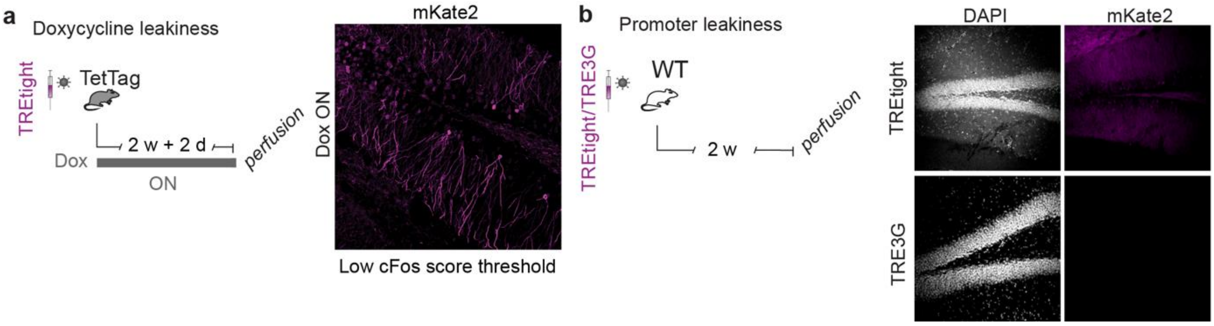


**Supplementary Figure 3. Optimization of cFos tagging in TetTag mice. a**, Doxycycline leakiness test. Mice were always ON Dox. They were injected with AAV9-TREtight-iChloC-mKate2 bilaterally in DG and were always in their home cage (HC). Few mKate2-expressing granule cells (GCs) could be detected. **b**, Promoter leakiness test. Wild-type mice that do not express the tetracycline-transactivator (tTA) were injected with either TREtight or TRE3G-promoter region constructs using AAVs. Right: mKate2 expression was observed in TREtight, but not with TRE3G promoter.

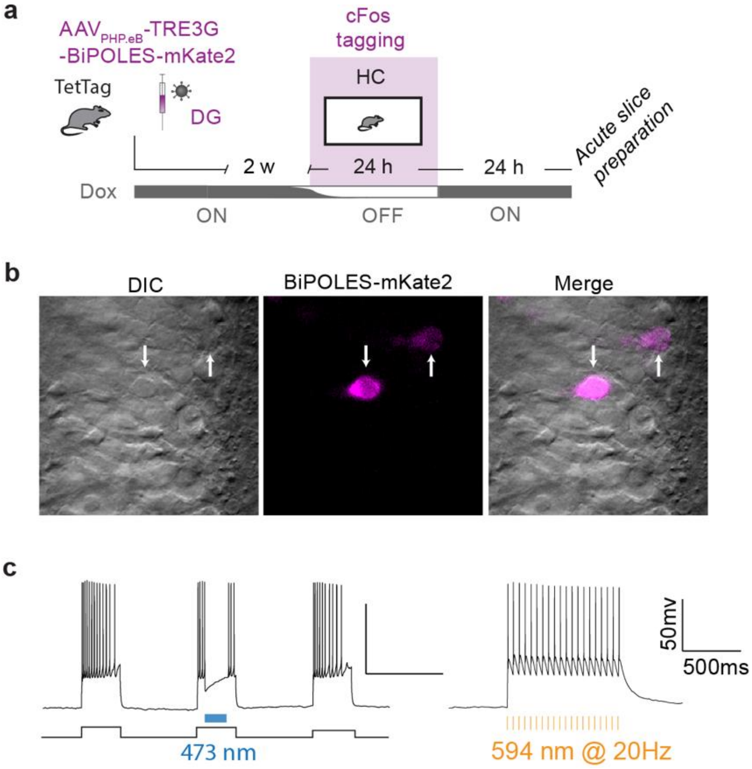


**Supplementary Figure 4. BiPOLES characterization in granule cells (GCs). a**, TetTag mice were injected with AAVPHP.eB-TRE3G-BiPOLES-mKate2. Mice were taken off Dox while they were in their home cage (HC). 24 h later, acute slices were prepared. **b,** Representative images showing the GC layer and BiPOLES-expressing cells. Left differential interference contras (DIC) image, middle: fluorescent image showing mKate2 (arrows) in the soma of GCs, right: overlay. **c,** Whole cell patch-clamp recordings from BiPOLES-expressing GCs. 473 nm light prevents action potentials (APs) induced by somatic 150 pA (500 ms) current injection. 594 nm light pulses reliably elicit APs.

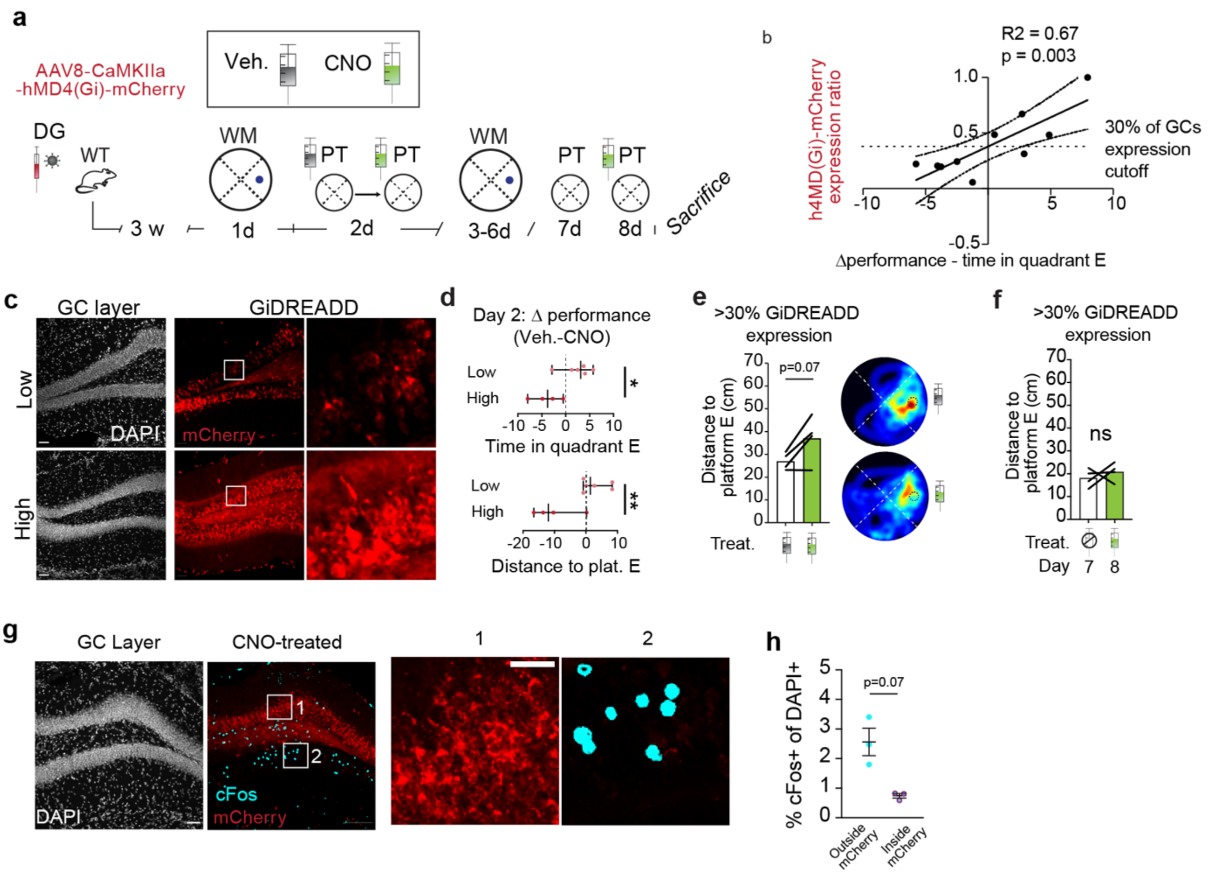


**Supplementary Figure 5. Chemogenetic silencing of dentate gyrus affects spatial memory recall. a**, Experimental design. Wild-type (WT) mice were injected with AAV8-CaMKIIa-hMD4(Gi)-mCherry bilaterally in DG (n = 10 mice). 3 weeks after injection, mice were trained in the water maze (WM) on day 1. On day 2, mice were tested for reference memory with two probe trials (PT, no platform). All mice were injected with 0.9% NaCl (vehicle) before the first PT and then with clozapine-N-oxide (CNO, 5mg/kg) before the second PT. Both injections were done intraperitoneally (ip) 40 min before each PT. Mice were trained further and tested again on day 7 for reference memory at the end of that session (no treatment) and the following day (8) 40 min after a CNO ip injection (without further training). Mice were sacrificed in different batches on different days with/without CNO application for immunohistochemistry. **b**, Expression of chemogenetic silencing receptors in DG strongly correlates with performance difference (with/without CNO) on day 2. Δ distance to platform E (CNO – Veh.) vs. hMD4i-mCherry expression ratio. The linear fit indicates improved performance for animals with very low GiDREADD expression, presumably reflecting the training effect of the first 'Atlantis' trial (without CNO): The target platform was raised at the end of the first probe trial. **c**, Representative confocal images showing hMD4(Gi)-mCherry expression in DG. Upper row: low Gi-DREADD-expressing mouse; lower row: high Gi-DREADD-expressing mouse. Left, DAPI. Middle, mCherry. Right, magnification from GCs expressing mCherry in the upper blade of the DG. Mice were separated into two groups based on their mCherry expression ratio: high > 0.5 (n = 4), low < 0.5 (n = 6). **d**, Mice in the low expression group are unaffected or show better performance on the second PT, while mice in the high expression group perform worse (***, $p = 0.0002$, unpaired t test). **e**, Chemogenetic silencing significantly impairs memory recall on day 2 in the high expression group (**, $p = 0.002$, paired t test). **f**, Spatial memory recall is not affected after they become experts. **g**, Example from a mouse where only the upper blade of DG was transduced with Gi-DREADD (red). No cFos expression (cyan) in granule cells of the upper blade (1), but strong cFos expression in GCs of the lower blade (2). This mouse received CNO before a PT and was sacrificed 1.5 h after. **h**, Data from three mice with uneven Gi-DREADD expression, comparing cFos expression outside and inside the (mCherry-labeled) Gi-DREADD expression area.

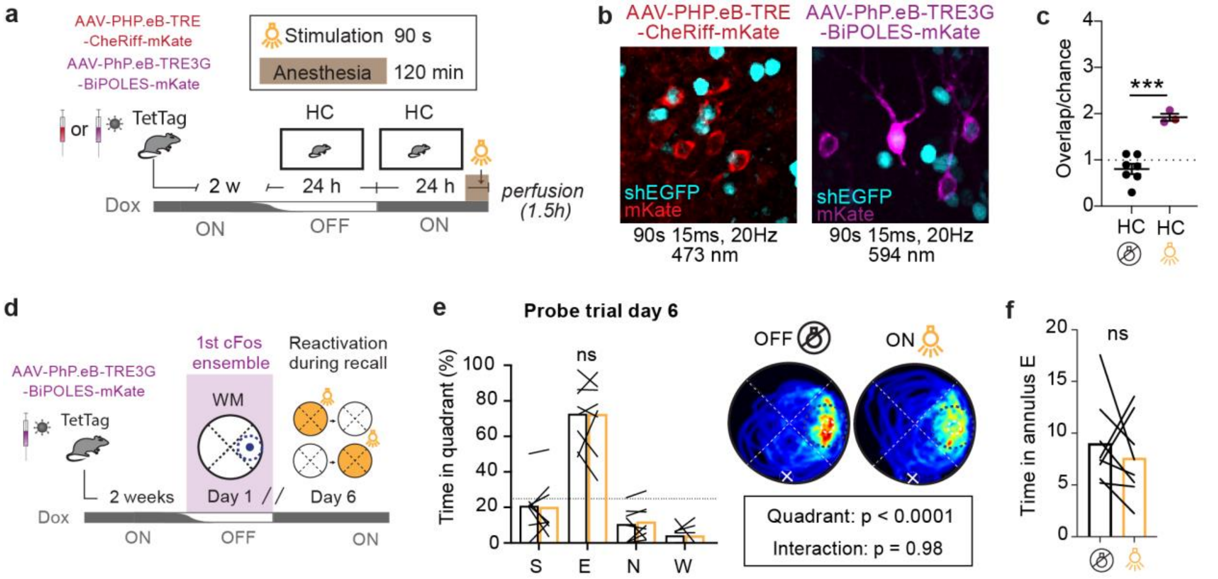


**Supplementary Figure 6. Effect of optogenetic reactivation of cFos$^+$ GCs on spatial memory recall. a,** Testing the effect of optogenetic depolarization in home-caged (HC) mice. Mice were taken off Dox for 24 h to express the optogenetic construct in cFos$^+$ cells. Two different channelrhodopsins were tested, CheRiff(red) and Chrimson (magenta), the depolarizing actuator in the BiPOLES construct. Animals were light-stimulated under anesthesia (473 nm for CheRiff, 594 for BiPOLES) and sacrificed 1.5 h later. **b, c**, Light stimulation increased the overlap between the first set of cFos$^+$ neurons (red/magenta) and shEGFP expression 24 h later (cyan). The difference to non-transduced animals (NT) was significant. **d**, Testing the effect of optogenetic reactivation of the cFos ensemble tagged on day 1 of WM training. All mice had two probe trials on day 6, one of which was performed under reactivation conditions (594 nm light pulses at 20 Hz). **e**, On day 6, performance in probe trials with optogenetic reactivation (orange bars) was not different from trials without reactivation (black bars). Overall performance was very good. **f**, Average distance to target platform was also not affected in trials with optogenetic reactivation (orange bar).
